# Local extinction of a montane, endemic grassland bird driven by landscape change across its global distribution

**DOI:** 10.1101/695536

**Authors:** Abhimanyu Lele, M. Arasumani, C. K. Vishnudas, Devcharan Jathanna, V. V. Robin

## Abstract

**Context:** Tropical montane habitats support high biodiversity, and are hotspots of endemism, with grasslands being integral components of many such landscapes. The montane grasslands of the Western Ghats have seen extensive land-use change over anthropogenic timescales. The factors influencing the ability of grassland-dependent species to persist in habitats experiencing loss and fragmentation, particularly in montane grasslands, are poorly known.

**Objectives:** We studied the relationship between the Nilgiri pipit Anthus nilghiriensis, a threatened endemic bird that typifies these montane grasslands, and its habitat, across most of its global distribution. We examined what habitat features make remnants viable habitat, which is necessary for effective management.

**Methods:** We conducted 663 surveys in 170 sites and used both single-season occupancy modelling and N-mixture modelling to account for processes influencing detection, presence, and abundance.

**Results:** Elevation had a positive influence on species presence, patch size had a moderate positive influence and patch isolation a moderate negative influence. Species abundance was positively influenced by elevation and characteristics related to habitat structure, and negatively influenced by the presence of invasive woody vegetation.

**Conclusions:** The strong effect of elevation on the highly range-restricted Nilgiri pipit makes it vulnerable to, and an indicator of, climate change. This highly range-restricted species is locally extinct at several locations and persists at low densities in remnants of recent fragmentation, suggesting an extinction debt. Our findings indicate a need to control and reverse the spread of exotic woody invasives to preserve the grasslands themselves and the specialist species dependent upon them.

## 1.0 Introduction

Tropical montane habitats are highly biodiverse, and harbour high endemicity (Ricketts et al., 2005; Dimitrov, Nogués-Bravo, & Scharff, 2012). They are also hotspots of extinction risk, due to the presence of threatened species with restricted distributions (Ricketts et al., 2005; Hoffmann et al., 2010). Montane specialists may also be threatened by climate change, which can trigger elevational range shifts (Stuhldreher & Fartmann, 2018). Where such shifts are limited by topography, species may face habitat decline and local extinctions (Parmesan, 2006; Forero-Medina, Joppa, & Pimm, 2011; Freeman, Scholer, Ruiz-Gutierrez, & Fitzpatrick, 2018). Habitat losses often also cause habitat fragmentation, which has experimentally been shown to have negative effects on biodiversity and species persistence over and above the effects of just habitat loss (Fahrig, 2003; Haddad et al., 2015). Declines in species abundances may occur after a significant time-lag following an environmental perturbation, creating a cryptic extinction debt, causing the effects of habitat disturbance to be underestimated (Tilman et al., 1994; Kuussaari et al., 2009; Haddad et al., 2015). Globally, extinction debt averages over 20% and may be as high as 75% of a local species assemblage (Haddad et al., 2015). The effects of climate change may interact with those of habitat loss and fragmentation (Fahrig, 2003), threatening montane habitats and the unique species assemblages they host.

The Western Ghats mountain range in southern India is a global biodiversity hotspot (Myers, Mittermeier, Mittermeier, & Kent, 2000) that includes locations of high extinction risk (Ricketts et al., 2005). The sky islands at the highest elevations of the Western Ghats host a naturally bi-phasic mosaic of evergreen forest and grassland known as the *shola* ecosystem. The montane grasslands of these Shola Sky Islands dominate the Western Ghats above 2000m (Das, Nagendra, Anand & Bunyan, 2015), and harbour unique species assemblages (Sankaran, 2009; Biju, Garg, Gururaja, Shouche, & Walujkar, 2014). As with other tropical grasslands, these are poorly studied, despite the presence of several endemic species, and others of conservation concern (Bond & Parr, 2010).

This grassland biome faces severe anthropogenic threats. Although the forests are celebrated for their biodiversity, historically, the ecological role of the grasslands has not been recognized, and they have been intensively exploited for the establishment of commercial plantations (Joshi et al., 2018). Many timber species thus introduced, including *Acacia mearnsii* (Black Wattle), *Eucalyptus* species, and *Pinus* species, have turned invasive (Arasumani et al., 2018; Joshi et al., 2018). Grassland loss to these species has been extensive, and is variously estimated at 83% overall (Sukumar, Suresh, & Ramesh, 1995), 66% in the Palani hills region (Arasumani et al., 2018) and 38% overall (Arasumani et al., 2019), depending on the spatial scale and temporal timeframe over which this change is measured. In addition to reducing habitat extent, the spread of exotic tree species has caused the grasslands, already a naturally patchy ecosystem (Robin, Gupta, Thatte & Ramakrishnan, 2015), to become further fragmented. These changes threaten the wildlife of the habitat, including endemic species such as the Nilgiri tahr *Nilgiritragus hylocrius* (Rice, 1984) and non-endemics that are supported by the grasslands (Sankaran, 2009). Effective conservation of these habitat specialists therefore requires understanding factors determining the persistence of native grassland-dependent species in the context of ongoing changes to the habitat.

The Nilgiri pipit, endemic to these montane grasslands, is an ideal case study to examine species persistence in this habitat. It is a locally common insectivore, resident in its breeding range, with no records of long-distance movement (Vinod, 2007). The small range of individual Nilgiri pipits and the presence of the species in small grassland patches including near human habitations make it a better indicator species for habitat change than the Nilgiri tahr, another species associated with this habitat, that can be impacted by hunting and other direct anthropogenic pressure (Mishra & Johnsingh, 1998).

Additionally, the effects of habitat characteristics on the distribution and abundance of the Nilgiri pipit are of wider interest in applied ecology: though features at both local and landscape scales have been shown to affect the presence and abundance of grassland birds, their habitat requirements in montane habitats have received little attention. At the local scale, changes to vegetation structure in native grasslands favours habitat generalists (Jacoboski, Paulsen, & Hartz, 2017). Microhabitat diversity within grasslands has been found to support a broader suite of grassland species (Diaz et al., 2014), and increased abundance of grassland-dependent species (Muchai, Lens, & Bennun, 2002; Azpiroz et al., 2012). More generally, structural changes in vegetation can affect species even in areas where native vegetation cover is high (Fischer & Lindenmayer, 2007). Features which create heterogeneity at small scales may be responsible for fragmentation at larger scales (Tews et al., 2004). Habitat heterogeneity in grasslands and montane habitats has received low attention (Tews et al., 2004). At the scale of a habitat fragment, area and isolation of grassland both affect grassland bird populations. Fragment area usually affects species’ occurrence; however, this effect may disappear beyond a certain threshold, which may be species-specific (Guttery et al., 2017). Local and landscape-level factors can have additive effects on occupancy (Reidy, Thompson, Amundson, & O’Donnell, 2016), and characteristics at different scales may act independently on occupancy and abundance (Blevins, 2011).

Finally, the Nilgiri pipit is itself a species of conservation concern. It is classified as vulnerable by the IUCN, due to its small and fragmented range, which is declining in both extent and quality (BirdLife International, 2018). Recent surveys have failed to detect the species across a significant portion of its historical range (Vinod, 2007; Robin, Vishnudas, & Ramakrishnan, 2014), suggesting that the contemporary range of the species is much smaller than expected. In this context, examining the habitat factors allowing the species to persist assumes greater importance. In this study we assessed factors driving patterns of distribution and abundance of the Nilgiri pipit across most of its known range. We used the single-season occupancy model of Mackenzie et al. (2002) to understand factors driving distribution, and the *N*-mixture model of Royle (2004) to understand grassland patch-specific variation in abundance.

## 2.0 Materials and methods

### 2.1 Study area

Robin et al. (2014) suggested that the Nilgiri pipit was restricted to areas above 1900 metres above sea level (a.s.l.). Conservatively, we limited our survey to grasslands above 1600m a.s.l., and also confined our study to areas in which verifiable contemporary records of the Nilgiri pipit exist. This region encompasses the two major high-altitude plateaux of the Western Ghats; the Nilgiris and the Anamalai-Palani Hills. Although the species has been reported elsewhere, photographic evidence or capture records for these locations, do not exist and intensive surveys across the smaller northern and southern grasslands have failed to detect the species (Robin & Sukumar, 2002; Robin et al., 2006; Sasikumar, Vishnudas, Raju, Vinayan, & Shebin, 2011).

**Figure 1.**
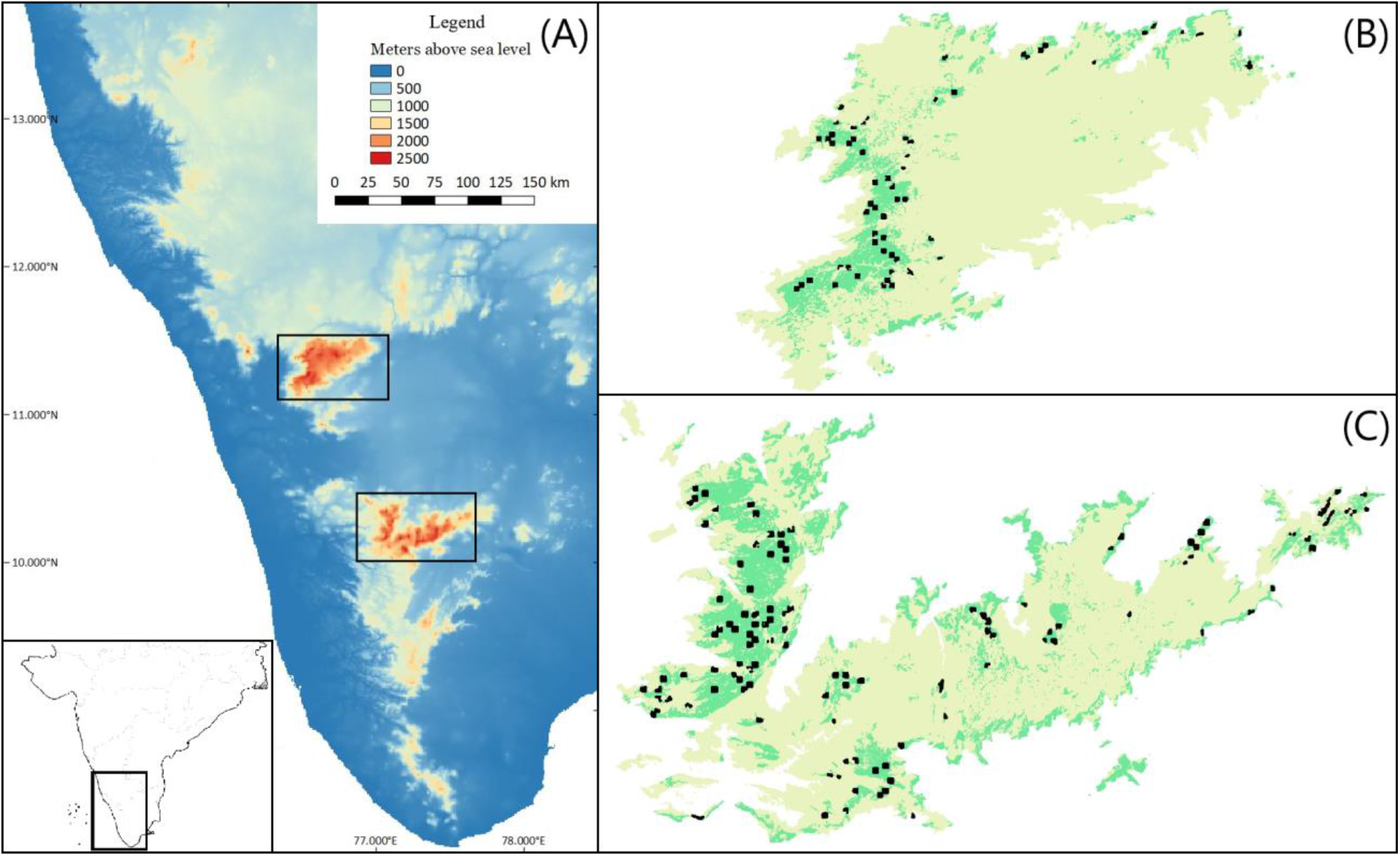
A: Map depicting the sky islands of the Western Ghats, and their position in the Indian subcontinent (inset). B-C: The grasslands (green) and sampling locations (black) in the Nilgiris (B) and the Anamalai-Palani Hills (C): areas in yellow are above the 1600m contour.

Within the above area, we mapped the extent of montane grasslands using Sentinel-2A imagery. We obtained imagery from the USGS Global Visualization Viewer (GloVis; https://glovis.usgs.gov/). Satellite imagery were obtained from the dry season (February 2017), when cloud cover was low. We removed atmospheric components such as dust particles, water vapour, and atmospheric temperatures in the satellite images by generating ground reflectance images using the Sen2Cor processor in SNAP v. 5.0.8 (ESA, 2017). We used a combination of supervised and unsupervised classification to map the montane grasslands (see supplementary methods for further detail). The overall accuracy of this classification, determined using 100 ground-truth GPS locations across the entire study area, was 96.5%, while the Kappa coefficient was 0.93 (following Congalton, 1991).

The final selected area represented 434.98km^2^, or 85%, of the 511km^2^ of grassland above 1600m in the Western Ghats. It consisted of 1,449 discrete grassland patches, varying in size from <1ha to 120,000ha. Grassland patches between 4 and 25ha were designated as sampling units, encompassing the range of our estimates for Nilgiri pipit home range (Vinod 2007, Vinod, U., personal communication, 2018). We placed patches between 1 and 4ha into clusters, if each patch was within a maximum of 200m (based on available information about pipit movement; Vinod, U., personal communication, 2018) away from at least one other, ensuring that slightly fragmented grasslands which were effectively larger than 4ha were not discarded. We determined the total area of each cluster, and discarded clusters or individual patches totalling less than 4ha as these were unlikely to support the species’ occurrence. We laid a 500m grid across all grassland patches larger than 25ha, designated each grid cell a separate patch, and removed all patches smaller than 4ha. We randomly selected 202 sites from the remaining 2,378 patches, and surveyed 170 of these (see Supplementary Methods for more details of survey design and site selection).

### 2.2 Survey methods

To accurately model both detection and occupancy, we visited each site multiple times (Max. visits = 4: 11 sites visited thrice, one site twice, one site once), with the duration of each visit set proportional to site area: species detection probability was thus equal across patches of variable size. We expended 2 minutes of survey effort per hectare, which we determined to be optimal based on reconnaissance surveys. Thus, surveys were between 8 and 50 minutes long. Surveys were conducted on foot; Nilgiri pipit counts were recorded based on visual and auditory detections strictly within the sampling site. Covariates were recorded at the site-level and the visit-level (Table 1: see Supplementary Methods for all covariates and definitions).

**Table 1:**
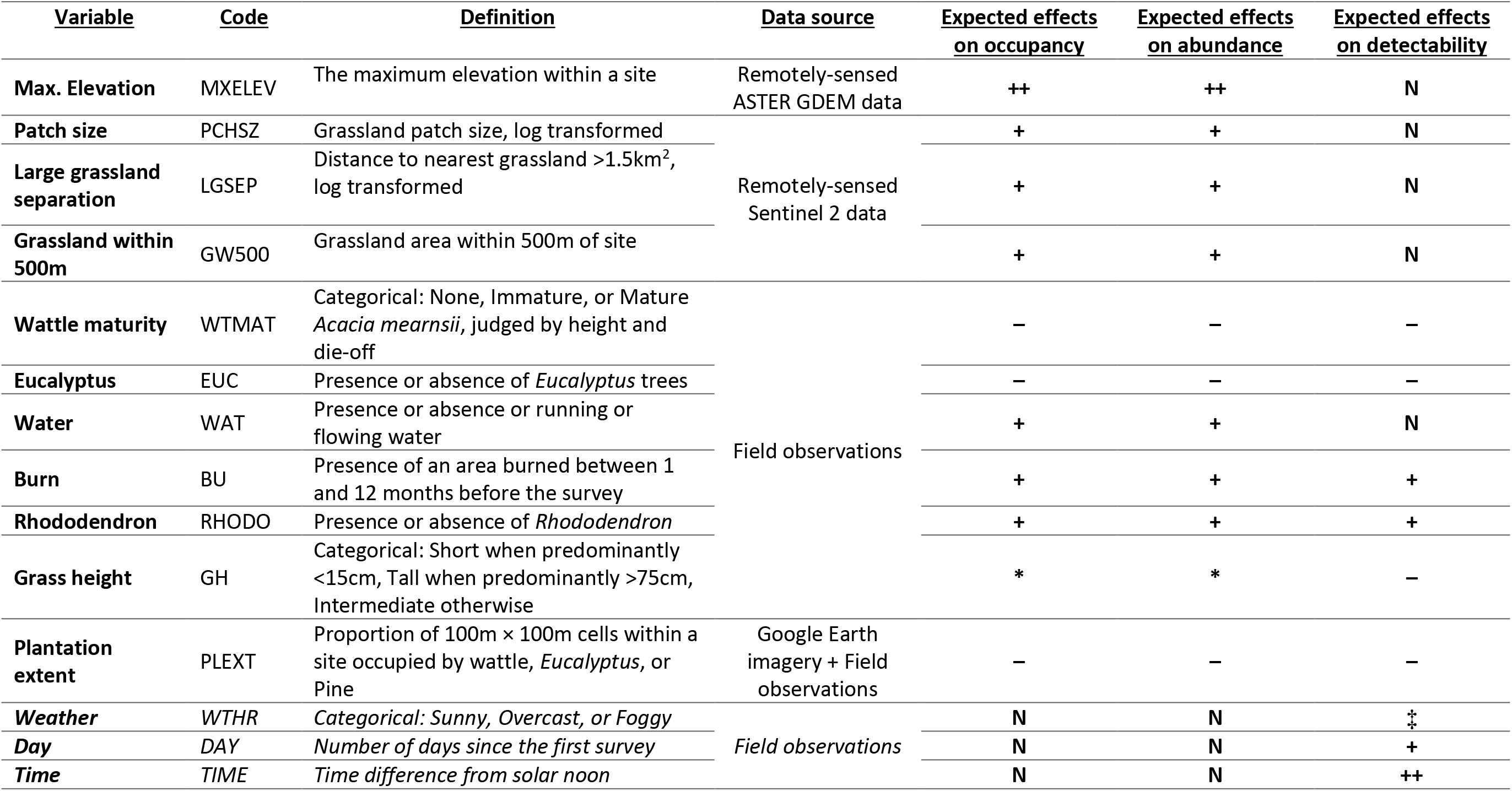
Description of variables used as predictors of Nilgiri pipit occupancy, abundance, and individual-level detectability. Visit level-variables are indicated in italics. Species-level detectability effects include individual-level detectability effects and abundance effects. ++ indicates a strongly positive expected effect; + indicates a positive expected effect; – indicates a negative effect: N indicates no expected effect. *For grass height, occupancy and abundance were expected to be highest for the intermediate category. ‡Detectability was expected to be low in foggy weather, higher in sunny weather, and highest in overcast weather. All expectations were derived from Vinod (2007; personal communication, 2018), Robin et al. (2014) and preliminary field surveys.

### 2.3 Statistical analysis

#### 2.3.1 Modelling Nilgiri pipit occupancy

Of the 22 site-level covariates, we eliminated some based on collinearity. Others were modelled only as covariates of detectability in our occupancy modelling (all details of variables in Supplementary Methods). Three visit-level variables were used to model detection probability of the species within sites.

Analyses were carried out using the package “unmarked” (Fiske and Chandler 2011) in the statistical software R (R Core team 2018). Counts were reduced to detections and non-detections for the occupancy analysis. To limit the number of models fit to the data, we carried out a two-step model fitting procedure. First, we used a general covariate structure to model the probability of occupancy and assessed support for covariates expected to affect detection of the species either directly, or indirectly by affecting abundance. We modelled additive effects of combinations of covariates, except in the case of Time and Weather, which we expected to interact in determining detectability. Model selection was carried out based on Akaike’s Information Criterion (AIC), and the best-fitting detection structures were then used to fit candidate occupancy covariate structures (following Doherty, White, and Burnham (2012)).

We tested 127 detection structures with an occupancy structure that included three covariates and used the best-fitting detection structures to fit 56 occupancy covariate structures. The best-fitting occupancy structures (AIC weight >0.02) are in Table 2: the best-fitting detection structures are in Supplementary Table 2. Since no single structure received overwhelming support, model-averaged predictions were used to derive response curves for each covariate (Burnham & Anderson, 2002). Since the final set of sites visited was non-random due to attrition of sites (see Supplementary Information for details), we did not attempt to estimate overall occupancy (proportion of area occupied) of Nilgiri pipits across the survey landscape.

**Table 2:** Estimated *β* coefficients for each predictor of occupancy from models with AIC weight ≥ 0.02. In each model presented below, detectability was modelled as a function of (Weather + Day + Grass height + Wattle maturity + Eucalyptus + Rhododendron + Water + Burn + Grassland within 500m).

| Occupancy structure | $\beta$<br>MXELEV | $\beta$<br>PCHSZ | $\beta$<br>LGSEP | $\beta$<br>WTMAT<br>Imm. Mat. | | $\beta$<br>EUC | $\beta$<br>PLEXT | $\beta$<br>RHODO | $\beta$<br>GH<br>Int. Tall | | AIC | ΔAIC | AIC Weight | Cum. AIC weight |
| --- | --- | --- | --- | --- | --- | --- | --- | --- | --- | --- | --- | --- | --- | --- |
| MXELEV+LGSEP | 9.01<br>$\pm 2.68$ | | -0.570<br>$\pm 0.266$ | | | | | | | | 537 | 0 | 0.173 | 0.173 |
| MXELEV+PCHSZ | 8.86<br>$\pm 2.43$ | 0.573<br>$\pm 0.251$ | | | | | | | | | 537 | 0.04 | 0.170 | 0.343 |
| MXELEV+LGSEP<br>+PLEXT | 9.55<br>$\pm 2.62$ | | -0.466<br>$\pm 0.277$ | | | | -1.37<br>$\pm 1.33$ | | | | 538 | 1.11 | 0.0992 | 0.442 |
| MXELEV+PCHSZ<br>+PLEXT | 9.39<br>$\pm 2.56$ | 0.464<br>$\pm 0.299$ | | | | | -0.939<br>$\pm 1.42$ | | | | 538 | 1.61 | 0.0772 | 0.520 |
| MXELEV+PLEXT | 10.8<br>$\pm 2.57$ | | | | | | -2.09<br>$\pm 1.21$ | | | | 539 | 1.79 | 0.0708 | 0.590 |
| MXELEV | 10.7<br>$\pm 2.83$ | | | | | | | | | | 539 | 2.05 | 0.0623 | 0.653 |
| MXELEV<br>+WATMAT | 13.0<br>$\pm 3.45$ | | | -2.45<br>$\pm 1.58$ | -2.15<br>$\pm 1.14$ | | | | | | 539 | 2.05 | 0.0623 | 0.715 |
| MXELEV+PLEXT<br>+GH | 9.34<br>$\pm 2.23$ | | | | | | -2.31<br>$\pm 1.07$ | | 1.78<br>$\pm 1.09$ | -0.44<br>$\pm 1.48$ | 539 | 2.45 | 0.0506 | 0.766 |
| MXELEV+PCHSZ<br>+RHODO+GH | 7.75<br>$\pm 2.69$ | 0.548<br>$\pm 0.222$ | | | | | | 0.123<br>$\pm 0.888$ | 1.61<br>$\pm 0.981$ | -0.15<br>$\pm 1.57$ | 540 | 3.51 | 0.0300 | 0.795 |
| MXELEV+PLEXT<br>+RHODO | 11.4<br>$\pm 3.38$ | | | | | | -2.21<br>$\pm 1.32$ | -0.348<br>$\pm 1.02$ | | | 540 | 3.67 | 0.0277 | 0.823 |
| MXELEV<br>+RHODO | 11.6<br>$\pm 3.99$ | | | | | | | -0.437<br>$\pm 1.13$ | | | 541 | 3.88 | 0.0249 | 0.848 |
| MXELEV+EUC | 10.4<br>$\pm 2.87$ | | | | | -0.314<br>$\pm 0.942$ | | | | | 541 | 3.94 | 0.0241 | 0.872 |
| MXELEV+LGSEP<br>+RHODO+GH | 7.71<br>$\pm 2.76$ | | -0.533<br>$\pm 0.239$ | | | | | 0.0134<br>$\pm 0.947$ | 1.49<br>$\pm 1.11$ | -1.09<br>$\pm 1.55$ | 541 | 4.04 | 0.0230 | 0.895 |
| MXELEV+LGSEP<br>+PLEXT+RHODO<br>+GH | 8.68<br>$\pm 3.17$ | | -0.385<br>$\pm 0.256$ | | | | -0.165<br>$\pm 1.22$ | -0.0461<br>$\pm 1.01$ | 1.53<br>$\pm 1.04$ | -0.696<br>$\pm 1.44$ | 541 | 4.26 | 0.0205 | 0.915 |

#### 2.3.2 Modelling Nilgiri pipit abundance

We used model-averaged occupancy predictions to estimate occupancy for each site. To avoid over-dispersion in site-specific abundance due to zero-inflation (Joseph, Elkin, Martin, & Possingham, 2009), only 112 sites with a predicted occupancy above 0.4, based on a clear threshold in a plot of estimated occupancy, were used for analysis using *N*-mixture models (Royle, 2004). A total of 22 detection covariate structures were tested with a global abundance covariate structure (see Supplementary Table 3 for selection statistics). We used the best-fitting covariate structure for detection to fit 91 covariate structures for abundance: six had an AIC weight of >0.02 (Table 3). Seven of the 11 independent variables appeared in structures with substantial support. We used log(site area) as an offset in all *N*-mixture models to control for site area, so that estimates of site-specific abundance λ could be interpreted as expected Nilgiri pipit density per hectare.

**Table 3:** Estimated *β* coefficients for each predictor of abundance from models with QAIC weight ≥ 0.02. In each model presented below, detectability was modelled as a function of (Weather + Day + Plantation cover).

| Abundance structure | $\beta$<br>MXELEV | $\beta$<br>WTMAT | | $\beta$<br>EUC | $\beta$<br>WAT | $\beta$<br>RHODO | $\beta$<br>GH | | $\beta$<br>BU | QAIC | ΔQAIC | QAIC Weight | Cum. QAIC weight |
| --- | --- | --- | --- | --- | --- | --- | --- | --- | --- | --- | --- | --- | --- |
|  |  | Imm. | Mat. |  |  |  | Int. | Tall |  |  |  |  |  |
| MXELEV+WTMAT<br>+EUC+WAT<br>+GH+BU | 1.424<br>$\pm 0.370$ | 0.313<br>$\pm 0.189$ | -0.434<br>$\pm 0.247$ | -0.377<br>$\pm 0.168$ | 0.504<br>$\pm 0.106$ | | 0.398<br>$\pm 0.193$ | -1.663<br>$\pm 1.010$ | 0.231<br>$\pm 0.122$ | 1018.63 | 0 | 0.298 | 0.298 |
| MXELEV+WTMAT<br>+EUC+WAT+RHODO+<br>GH+BU | 1.303<br>$\pm 0.383$ | 0.310<br>$\pm 0.188$ | -0.390<br>$\pm 0.247$ | -0.365<br>$\pm 0.168$ | 0.464<br>$\pm 0.110$ | 0.192<br>$\pm 0.147$ | 0.371<br>$\pm 0.195$ | -1.661<br>$\pm 1.007$ | 0.192<br>$\pm 0.125$ | 1018.92 | 0.281 | 0.259 | 0.557 |
| MXELEV+WTMAT<br>+EUC+WAT<br>+RHODO+GH | 1.310<br>$\pm 0.386$ | 0.356<br>$\pm 0.187$ | -0.332<br>$\pm 0.246$ | -0.367<br>$\pm 0.169$ | 0.474<br>$\pm 0.110$ | 0.241<br>$\pm 0.143$ | 0.303<br>$\pm 0.191$ | -1.713<br>$\pm 1.006$ | | 1019.26 | 0.630 | 0.218 | 0.775 |
| MXELEV+WTMAT<br>+ WAT+RHODO<br>+GH+BU | 1.24<br>$\pm 0.382$ | 0.374<br>$\pm 0.184$ | -0.339<br>$\pm 0.247$ | | 0.471<br>$\pm 0.111$ | 0.211<br>$\pm 0.148$ | 0.410<br>$\pm 0.197$ | -1.725<br>$\pm 1.007$ | 0.195<br>$\pm 0.126$ | 1021.68 | 3.046 | 0.065 | 0.840 |
| **MXELEV+WTMAT**<br>**+WAT+GH+BU** | 1.380<br>$\pm 0.369$ | 0.381<br>$\pm 0.185$ | -0.377<br>$\pm 0.247$ | | 0.511<br>$\pm 0.107$ | | 0.436<br>$\pm 0.196$ | -1.750<br>$\pm 1.018$ | 0.234<br>$\pm 0.123$ | 1021.72 | 3.083 | 0.064 | 0.904 |
| **MXELEV+WTMAT**<br>**+WAT+RHODO+GH** | 1.246<br>$\pm 0.385$ | 0.412<br>$\pm 0.184$ | -0.287<br>$\pm 0.246$ | | 0.481<br>$\pm 0.111$ | 0.258<br>$\pm 0.144$ | 0.344<br>$\pm 0.193$ | -1.744<br>$\pm 0.988$ | | 1022.07 | 3.434 | 0.054 | 0.957 |

## 3.0 Results

### 3.1 Nilgiri pipit occupancy

We detected the Nilgiri pipit in 109 of 170 sites (naïve occupancy = 0.641). Adequate model fit (*P* = 0.8392) was indicated by the χ^2^ goodness-of-fit test from 5,000 parametric bootstrap simulations on the top model (MacKenzie & Bailey, 2004), for the subset of sites visited four times (*N* = 156); we excluded sites visited thrice, twice and once, as sample sizes were too low to treat these as separate cohorts. Model-averaged results show that maximum site elevation had a strong positive effect on occupancy, while patch isolation and patch area (fit only in separate models, as they were strongly correlated) had moderate negative and positive correlations with occupancy, respectively (Figure 2). All 14 best-fitting models included maximum site elevation as a covariate, while the top four included either patch isolation or grassland patch area. All but one (quantity of grassland within 500m of the site) of the putative occupancy predictors appeared in one or more model with substantial support. Models comprising combinations of isolation, area, and elevation had comparable AIC weights to those including additional covariates: however, each of these other covariates had extremely small effects on occupancy.

**Figure 2.**
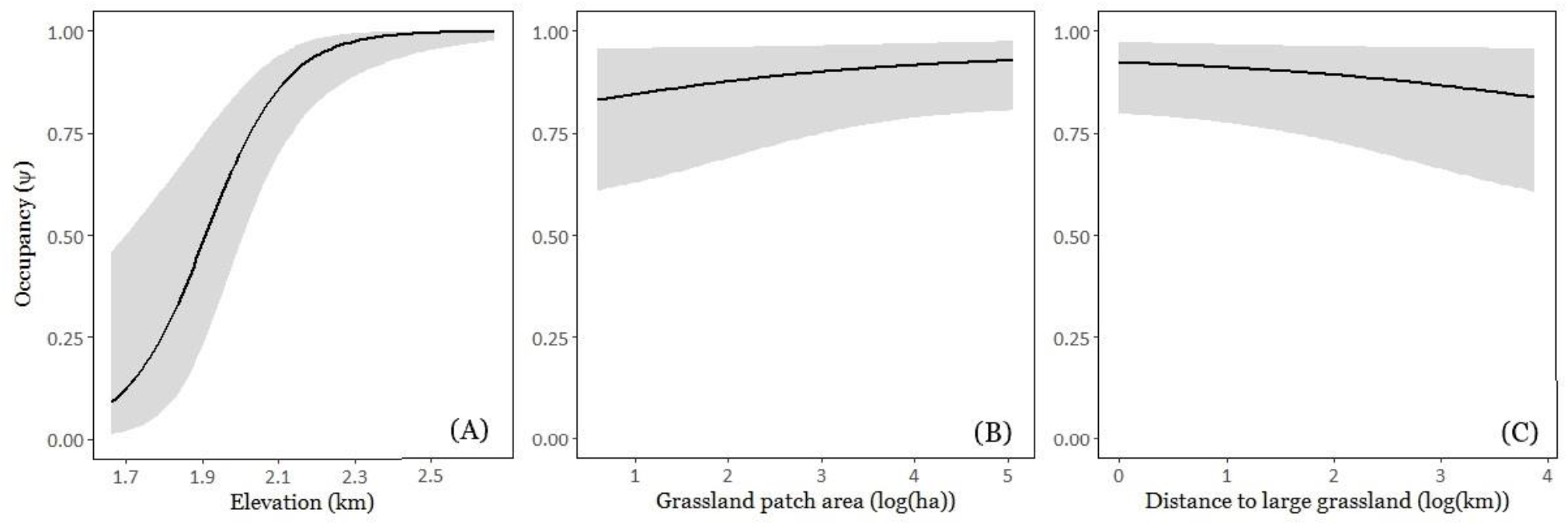
A-C: Model-averaged predicted occupancy of the Nilgiri pipit in response to the three covariates with the largest effects; maximum elevation within a site (A), log grassland patch area (B), and log distance to the nearest grassland larger than 1.5 km^2^ (C). All other variables are set to mean values. Predicted probability of occupancy is plotted over the observed range of values of each predictor.

### 3.2 Nilgiri pipit abundance

In the 144 sites with ψ > 0.4, we detected 0 to 14 individuals (mean = 3.76). Despite excluding sites with a low probability of occurrence, the *χ*^2^ goodness-of-fit test based on 5,000 parametric bootstrap simulations (MacKenzie & Bailey, 2004) indicated over-dispersion of latent abundances relative to the model (*ĉ*=1.93). Estimated *ĉ* was therefore used to derive QAIC values for model selection, and to adjust estimated variances. Model-averaged results showed that elevation had a strongly positive effect on bird density: the predicted density at the maximum sampled elevation was more than twice that at the lowest elevation. The best-fitting six models all included elevation, grass height, wattle maturity, and water, while *Rhododendron, Eucalyptus*, and recent burns appeared in three of the four best-fitting models. *Eucalyptus* negatively affected density, while the presence of water positively affected density. The presence of both *Rhododendron* and burned area had marginal positive effects on density. Grassland patch size and isolation, and plantation extent, were not part of models that had any substantial support.

**Figure 3.**
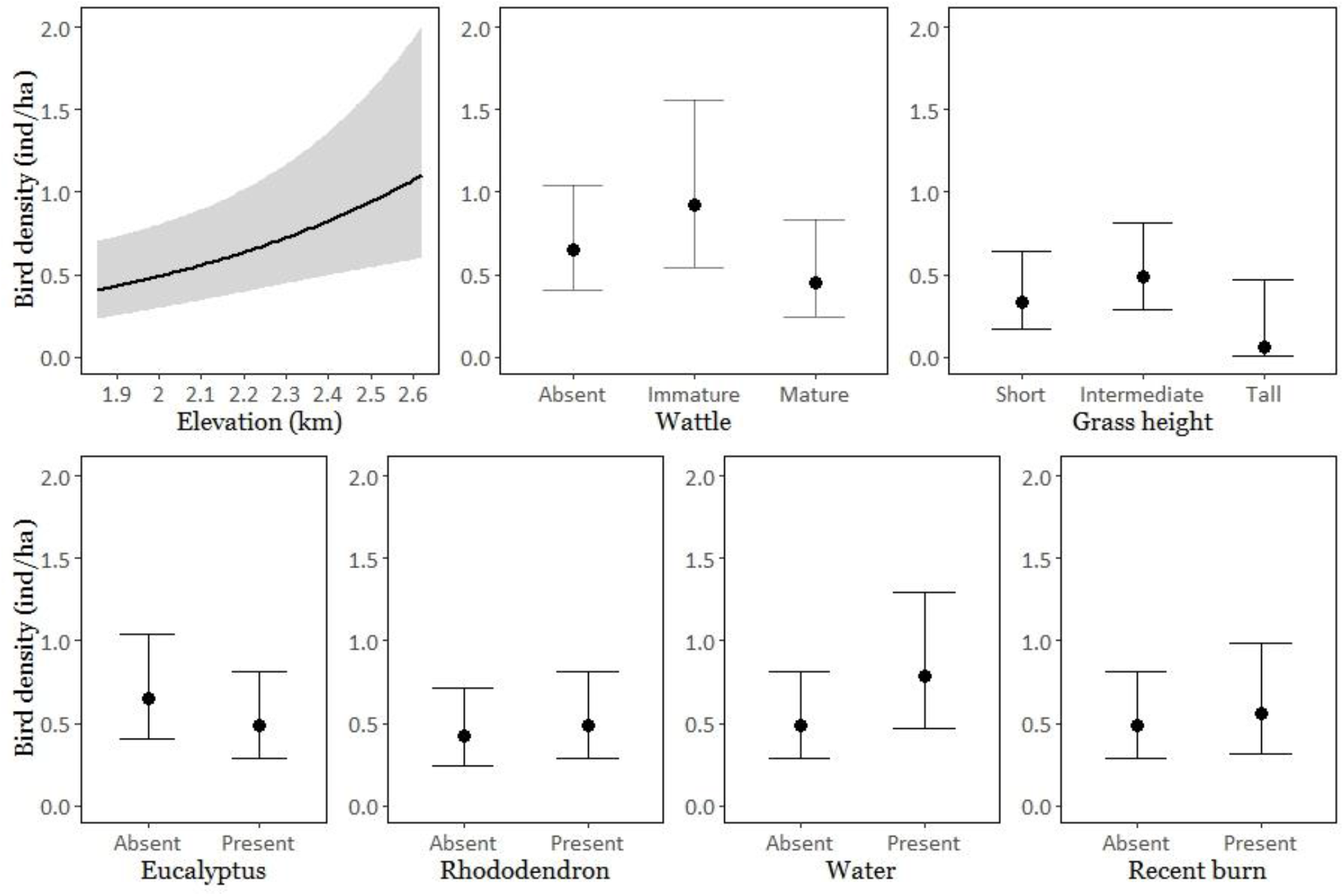
A-G: Model-averaged predicted density (per hectare) of the Nilgiri pipit in response to all covariates appearing in models with AIC weight >0.01; maximum elevation within a site (A), maturity of wattle (B), grass height (C), presence of *Eucalyptus* (D), presence of *Rhododendron* (D), presence of water (F), and presence of a recently burned area (G). All other variables are set to mean or modal values for continuous and categorical covariates, respectively. Predicted density is plotted over the observed range of values of each predictor.

## 4.0 Discussion

We found that maximum site elevation, grassland patch size, and distance to the nearest large grassland were the only covariates that had substantial effects on pipit occupancy. In contrast, abundance was shaped by maximum site elevation in combination with many site-level habitat characteristics, each of which had a substantial effect on predicted abundance.

Nilgiri Pipit occupancy and abundance both have strong relationships with elevation. Only sites above 1800m have high probabilities of occupancy. Thus, the Nilgiri pipit is likely to be the most elevationally restricted bird species in the Indian subcontinent south of the Himalaya. The only detection of Nilgiri pipits below 1700m was at the southwestern extremity of the Anamalai plateau, with anomalously high exposure to the southwest monsoon and microclimate consistent with much higher areas elsewhere. This relationship between occurrence and elevation is consistent with previous research indicating that montane grasslands become the dominant biome only above 2000m (Das et al., 2015), and is similar to the distribution patterns of other montane flora including *Rhododendron* (Giriraj et al., 2008) and fauna (Yandow, Chalfoun, & Doak, 2015; Mizel, Schmidt, Mcintyre, & Roland, 2016) including habitat-specialist birds (Watson, 2003) like the *Sholicola* (Robin & Sukumar, 2002). This sensitivity to elevation makes the Nilgiri Pipit extremely vulnerable to climate change: it is likely to undergo substantial range contraction due to anthropogenic global warming (Parmesan, 2006; Sekercioglu, Schneider, Fay, & Loarie, 2008), given the distribution of area with respect to elevation in the two plateaux (Table 4).

**Table 4:** Cumulative available grassland above successive 100m contours in the Western Ghats. The total available grassland declines rapidly with elevation, implying that the range of the Nilgiri pipit would decline rapidly if it is forced upwards by climate change.

| **Elevation a.s.l.**<br>**(m)** | 1600 | 1700 | 1800 | 1900 | 2000 | 2100 | 2200 | 2300 | 2400 | 2500 |
| --- | --- | --- | --- | --- | --- | --- | --- | --- | --- | --- |
| **Cumulative**<br>**area of**<br>**Grassland**<br>**(km<sup>2</sup>) above**<br>**each**<br>**elevation**<br>**contour** | 511.1 | 436.6 | 378.6 | 329.9 | 278.0 | 217.6 | 150.0 | 83.6 | 34.9 | 5.1 |

Any loss of habitat for restricted-range species such as the Nilgiri pipit can be severely detrimental. Our sampling did not include patches of dense, mature, plantations of the invasive woody species, as these habitats are not viable for grassland specialists. Within the sampled grasslands, the presence of these invasives did not affect the occupancy of the Nilgiri pipit, but negatively affected abundance. The exception was the patch condition “immature wattle”, which had a higher predicted abundance than either “mature wattle” or “no wattle”. We suggest two possible explanations for this anomaly. First, areas with immature wattle but without mature wattle largely (19 of 28 sites) occur within the two largest grassland patches, Eravikulam and Mukurthi National Parks, which have high Nilgiri pipit densities, and where management practices include removal of mature wattle. Second, immature wattle may temporarily contribute to habitat heterogeneity (a factor that increases abundance, as discussed below) without substantially degrading habitat quality. However, due to wattle’s rapid growth, this effect is likely to be transient.

Our findings suggest that while Nilgiri pipit presence may be constrained by habitat availability, its abundance is shaped by local habitat quality. Woody monocultures are driving declines in abundance: such declines may cause functional extinction even in areas where the species is present (Dirzo et al., 2014). Furthermore, woody invasives are replacing grassland over longer timescales (Arasumani et al., 2018), and thereby are likely to also constrain occupancy. Several studies have found detrimental impacts of spreading woody exotic species on grassland avifauna, including in the Brazilian Pampas (Azpiroz et al., 2012; Jacoboski et al., 2017) and South African highland grasslands (Allan, Harrison, Navarro, Van Wilgen, & Thompson, 1997), suggesting that such a spread is a widespread phenomenon globally, requiring broader attention.

It is probable that low-density populations in areas affected by invasive species are non-viable. In grasslands that are remnants of century-old habitat loss in the eastern Nilgiris (Joshi et al., 2018) we found a complete absence of Nilgiri pipits, but grassland remnants in the Palani Hills, invaded and fragmented severely since 1973 (Arasumani et al., 2018) supported low-density populations. We conclude that the eastern Nilgiris have experienced local extinction, as historical records of the species exist from that region (Robin et al., 2014). Furthermore, high-altitude habitat specialists are often strongly affected by patch area (Watson, 2003), but the Nilgiri pipit showed only a weak correlation with patch size and isolation, which is unexpected for a poor disperser (Rosenzweig, 1995), such as the Nilgiri pipit (Vinod, U., personal communication, 2018). These findings may represent a substantial extinct debt in grassland recently affected by invasive species in the Palani hills and the southern Anamalais. The effects of local extinctions on population structure and viability is likely to be stronger in non-avian grassland endemics, since birds have greater dispersal abilities (Watson, 2003), particularly for long-lived species that are likely to have greater extinction debt (Krauss et al., 2010).

The abundance of the Nilgiri pipit showed a strong positive correlation with intermediate or mixed grass height: such a preference for specific grass height has also been documented for a suite of avian species in the Brazilian pampas (Jacoboski et al., 2017). We found moderate positive correlations with other variables contributing to local habitat heterogeneity and vegetation structure, a correlation supported by the behavioural observations of Vinod (2007), who found that Nilgiri pipits preferred marshy habitat with tall grass for nesting, and more open habitat for feeding. The relationship between habitat heterogeneity and grassland birds is complex, with both positive and negative responses documented (Wiens & Rotenberry, 1981; Pavlacky, Possingham, & Goldizen, 2015). While further investigation of the functional effects of structural complexity in the montane grasslands of the Western Ghats is merited, our findings show a broad dependence on natural habitat heterogeneity within this habitat.

Our findings have several implications for land management. The contiguous grassland within Eravikulam and Grasshills National Parks is the only remaining patch with high pipit abundance without large areas lost to invasive species: its continued protection is therefore of critical importance. The grassland in and around Mukurthi National Park, containing the only other high-density population, is rapidly shrinking due to invasion (Arasumani et al., 2019), largely by black wattle, and requires urgent management attention. Conversely, many grasslands in which we suspect high potential for extinction are outside protected areas, and have already been severely fragmented. These require a different management strategy targeting connectivity and restoration: such strategies have already been explored at local scales (Mudappa, D., & Shankar Raman, T. R., personal communication, 2018; Stewart, R., personal communication, 2018)

We found that patch-level habitat quality had a strong effect on abundance: such a pattern has also been found in other highland avifauna (Allan et al., 1997; Watson, 2003). Thus, conservation efforts must focus on maintaining habitat quality over and above simply preserving grassland. Frequent fire is thought to positively affect grassland avian species richness (Pons, Lambert, Rigolot, & Prodon, 2003). We did not find a strong relationship between recent burns and Nilgiri pipit density: further systematic study is required to draw any conclusions about the role of fire in management in this landscape. At a local scale, the presence of woody invasives reduces Nilgiri pipit abundance, while at the landscape scale, the spread of the same woody invasives shapes grassland patch size and isolation (Arasumani et al., 2019), which drive Nilgiri pipit occupancy. We emphasise that our study was limited to the remnant grasslands, and our findings therefore underestimate the detrimental effects of invasive vegetation, as completely wooded habitats that were grasslands earlier were not sampled since these do not have any grasslands or pipits. Controlling the spread of invasive tree species is a matter of urgent conservation attention.

## 5.0 Conclusions

We demonstrate that elevation shapes both occupancy and abundance of this montane specialist. Furthermore, our study demonstrates that woody invasives are constraining occupancy via rapid grassland loss at the landscape level, while also degrading habitat quality at the local level. Our research also indicates local extinctions in large parts of the species’ range, and probable extinction debt in other parts. This study underscores the urgent need for conservation actions targeted at the poorly known montane grasslands and the specialist species dependent upon it.

## 7.0 Acknowledgements

We thank the Forest Departments of Kerala and Tamil Nadu for permits (WL5(A)/12260/2017, WL10-411/2017). We thank Forest Department officers P. G. Krishnan and R. Lakshmi for discussions and support to the project. We thank V. Joshi and A. Aravind for assistance with data collection; S. Ray and D. Khan for field assistance; V. Godwin, R. Pilakandy, R. Bhalla, K. Shanker, and D. Mudappa, for support during fieldwork; V. Godwin, V. Joshi, U. Vinod, and M. Bunyan, for discussions and critical comments on the manuscript. This study was supported by National Geographic Society grant WW-186EC-17 to AL, a grant from the Duleep Matthai Nature Conservation Trust to VVR and CKV, and IISER Tirupati funding to VVR. Revisions of this manuscript were made after AL started graduate school at the University of Chicago, and we thank them for their support. This research was carried in compliance with all relevant institutional norms and Indian laws.

## 8.0 Author contributions

VVR, CKV, DJ, and AL conceptualized and designed the study. AM collated and analysed remotely sensed data. AL and CKV collected field data. AL analysed field data with help from DJ and VVR. AL wrote the manuscript with input from all authors.

## 10.0 Supplementary methods

### 10.1 Administrative regions examined in the study

Our study area fell within the following administrative regions. In the Nilgiris, our surveys were conducted within Mukurthi National Park, Nilgiri South Forest Division, and Nilgiri North Forest Division, all units of the Tamil Nadu Forest Department. In the Anamalai-Palani Hills region, our surveys were carried out in Grasshills National Park, Kodaikanal Wildlife Sanctuary, and Dindigul Forest Division (under the Tamil Nadu Forest Department), and Eravikulam National Park and Munnar Wildlife Division under the Kerala Wildlife Department. A few survey locations were located within private land, in which case permission was obtained from the owners to conduct surveys therein.

### 10.2 Mapping grasslands using remote sensing

We used Sentinel - 2A data for montane grassland mapping. Satellite images were downloaded from USGS Global Visualization Viewer (GloVis; https://glovis.usgs.gov/): the images acquired were from February and March 2017. We selected images with less than 1% cloud cover from GloVis and acquired them for further processing. Nine Sentinel 2A scenes covered our study area. We mosaicked all the satellite images using SNAP v. 5.0.8. and performed a geometric correction to improve geolocation of each pixel: the root mean square error was less than half a pixel. We corrected for atmospheric distortion by removing atmospheric components (e.g. water vapour column, dust particles and aerosol) using the Sen2cor processor in SNAP v5.0.8. We used four bands (Band 2 – 192.4 nm, Band 39 - 559.8, Band 4 - 664.6 and Band 8 – 832.8) with 10m spatial resolution for image classification. Our study area was divided into 5 km^2^ grid cells for training sample collection and ground truth verification. We collected 223 training samples between April 2017 and September 2017 across the study area for image classification. We interpreted the nine Sentinel 2A scenes using the hybrid classification approach, which combines digital supervised classification, unsupervised classification, and topography (Arasumani et al. 2019). The accuracy of the classified map was calculated using ground truth GPS points. We created 100 random points on the map and visited each of them to evaluate the accuracy of our classification. Overall accuracy was found to be 96.5%, and the kappa coefficient was 0.93 (following Congalton, 1991).

### 10.2 Sampling sites and site selection

Of the 3012 grassland patches which remained after small patches had been removed from our sampling frame (*sensu* Williams, Nichols, & Conroy, (2002)), 300 were randomly selected to be surveyed for detection/non-detection of Nilgiri pipits. Of these 300, areas containing 69 sites were removed because they were determined by the relevant government agency to be unsafe due to armed militant activity, or for other reasons of accessibility. The remaining study landscape included 234 sites. Of these 234, 8 sites were removed due to misclassification: they were found to be *Eucalyptus* plantation or scrub forest, and in 3 cases, to be under water, because they occurred on the banks of a reservoir in which the water level had risen between the point at which classification was carried out, and the time at which the surveys were conducted. A further 21 sites could not be sampled as they were directly adjacent to other (sampled) sites, and Nilgiri pipit presence in them could not be assumed to be independent of all other sites. Thus, the total area of grassland sampled was 434.98km^2^, which contained 202 sites that fit our criteria for sampling (accurate habitat classification, and independence from other sites. Of these 202, 170, or 84%, were sampled: 32 sites were found to be physically inaccessible due to the topography of the landscape or the ownership of the land.

### 10.3 Covariates

Landscape-level covariates were generated from GIS data. Site-level covariates were generated either from GIS data or field observations. Visit-level covariates were measured in the field. A full list of covariates is provided below. Variables used in the final analyses are in **boldface** font. Of the 22 site-level independent variables, we eliminated some variables based on collinearity. Five variables were eliminated because there was insufficient variation in these across our sites. Any other covariates displaying moderate collinearity were not included within the same model. Of the 11 remaining covariates, two were expected to affect abundance but not occupancy, and some were used only as detection covariates in modelling pipit occupancy.

**Supplementary Table 1:**
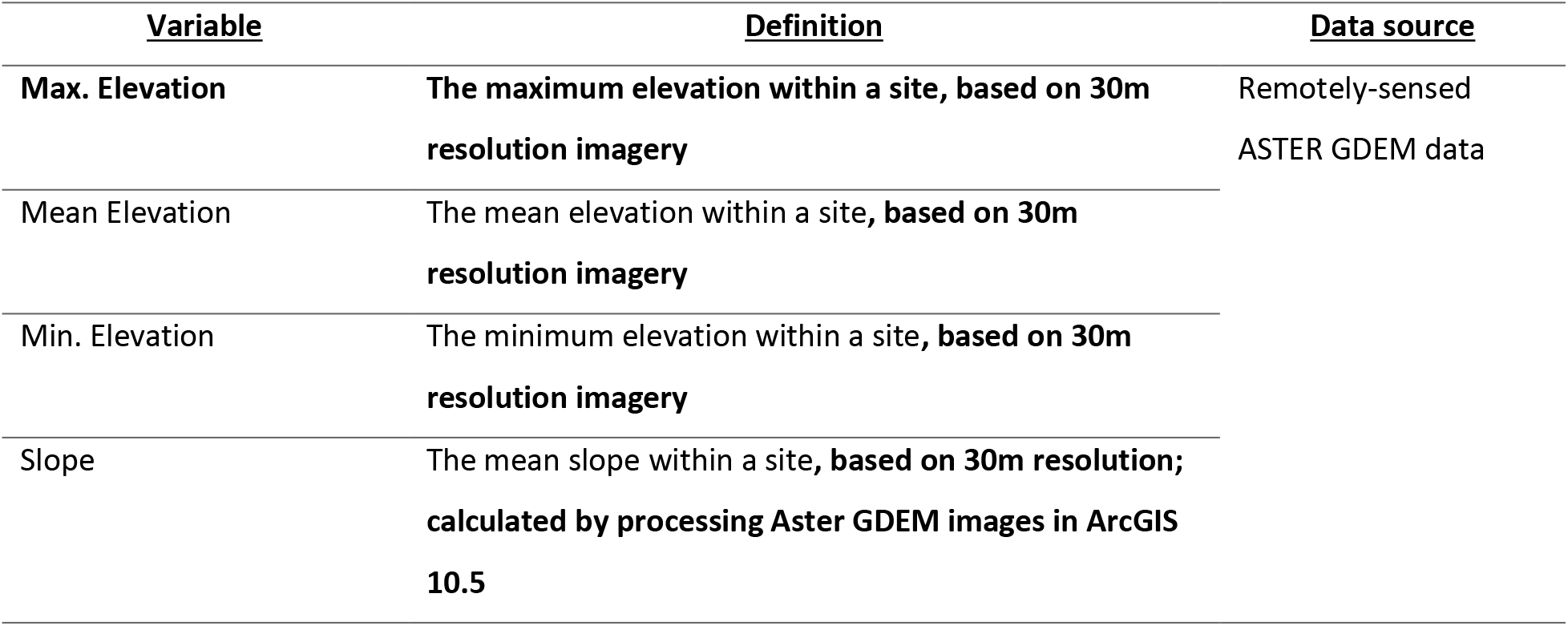
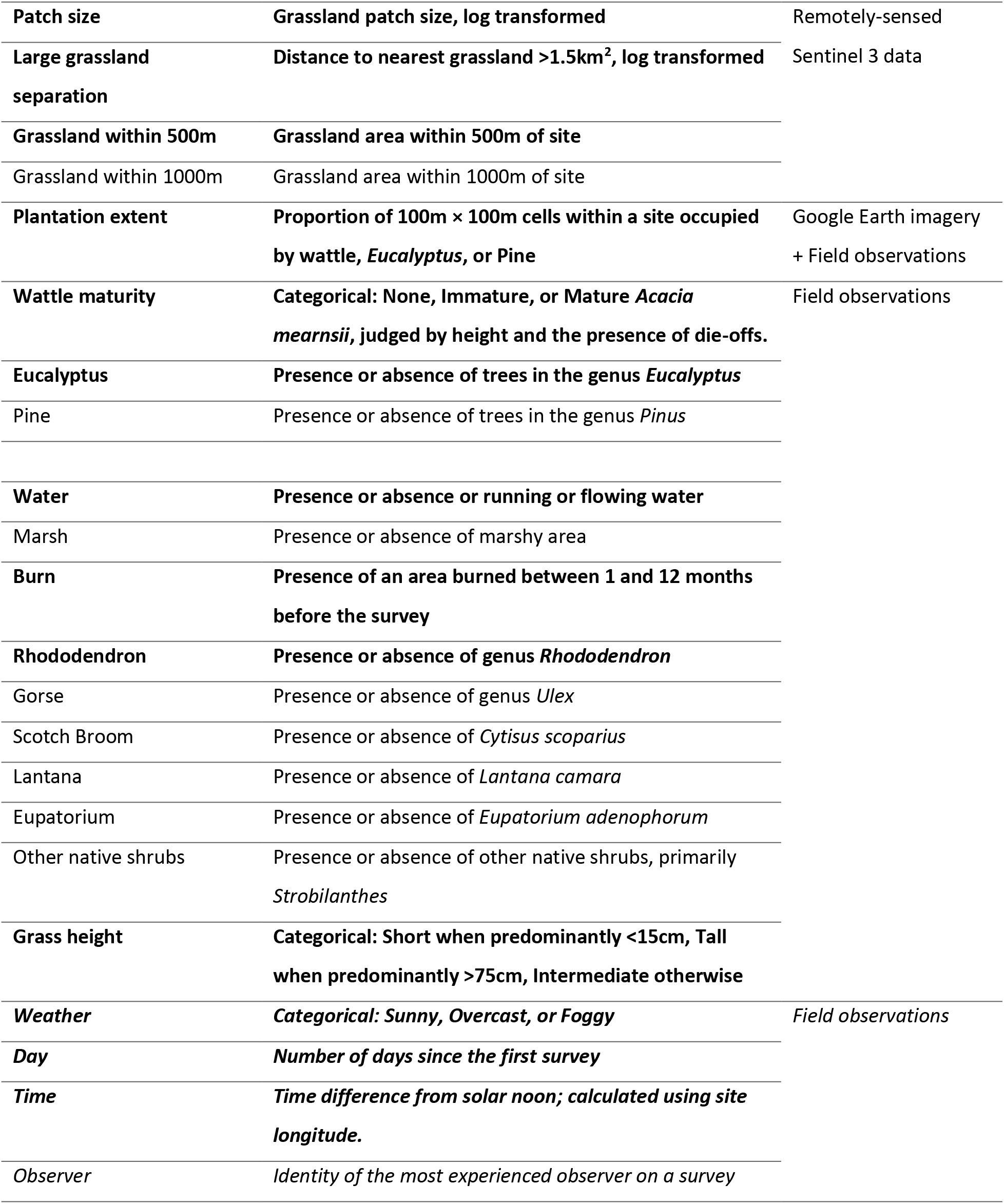
All covariates

Covariates included the presence or absence of the following categories of vegetation; black wattle (*Acacia mearnsii*), *Eucalyptus*, pine (genus *Pinus*), *Lantana camara, Eupatorium adenophorum* (also known as *Ageratina adenophora*), rhododendron (genus *Rhododendron*), and other native shrubs, including *Strobilanthes spp*. The first five of these vegetation types are common invasive exotics in the high-altitude grassland landscape; the other two covariates represent the other significant types of vegetation found within the grassland. Data for *A. mearnsii* were divided into the categories “Mature” and “Immature”, as each category was expected a priori to affect pipits differently and these also seemed to accurately reflect variation on the ground,; several sites had experienced recent invasion from *A. mearnsii* and had not yet been substantially altered by this invasion. Furthermore, black wattle trees experience die-offs beyond a certain size, which are easily identifiable in the field, providing a good proxy for the maturity and hence density of a stand of wattle.

The threshold of 1.5km^2^ for defining a “large grassland” was based on observations which found that virtually all pipit populations which appeared large and healthy, and where pipits were regularly detected in large numbers, were in patches larger than 1.5km^2^.

The predominant height of the grass within a site was a categorical variable with three level. Grass was characterized as short when it was shorter than 15cm, tall when it was taller than 75cm, and intermediate in all other cases. This categorization was based on observations of different microhabitats. Cliffs, heavily grazed areas, or recently burned areas had extremely short grass; marshy areas and scrub habitat found up to approximately 1800m had very tall grass; the ‘intermediate’ category typically included native montane grasslands and more mixed grassland.

Visit-level covariates included the weather, date, time of day, and identity of the most experienced observer. Weather was recorded as a categorical variable with three levels; sunny, overcast, and foggy, as these types of weather had clear effects on detection. Data were not collected during rain. Observer identity was recorded to assess potential differences in detection efficacy between observers. The number of days since the first survey was recorded as a proxy for season, as seasonal variation in the Nilgiri pipit’s behaviour was known (Vinod 2007) and needed to be controlled for. Similarly, the diurnal activity of the Nilgiri pipit was known to have a bimodal pattern with peaks soon after sunrise and shortly before sunset: thus, we transformed time of day to time away from solar noon for analysis.

Gorse, Scotch Broom, Pine, and *Lantana* were eliminated as variables because they were found to be present in <10% of the sites surveyed. *Eupatorium* and the “Other native shrubs” variable were eliminated because they were found in >95% of the sites surveyed.

### 10.4 Estimated effects of all putative covariates on occupancy

**Supplementary Figure 1.**
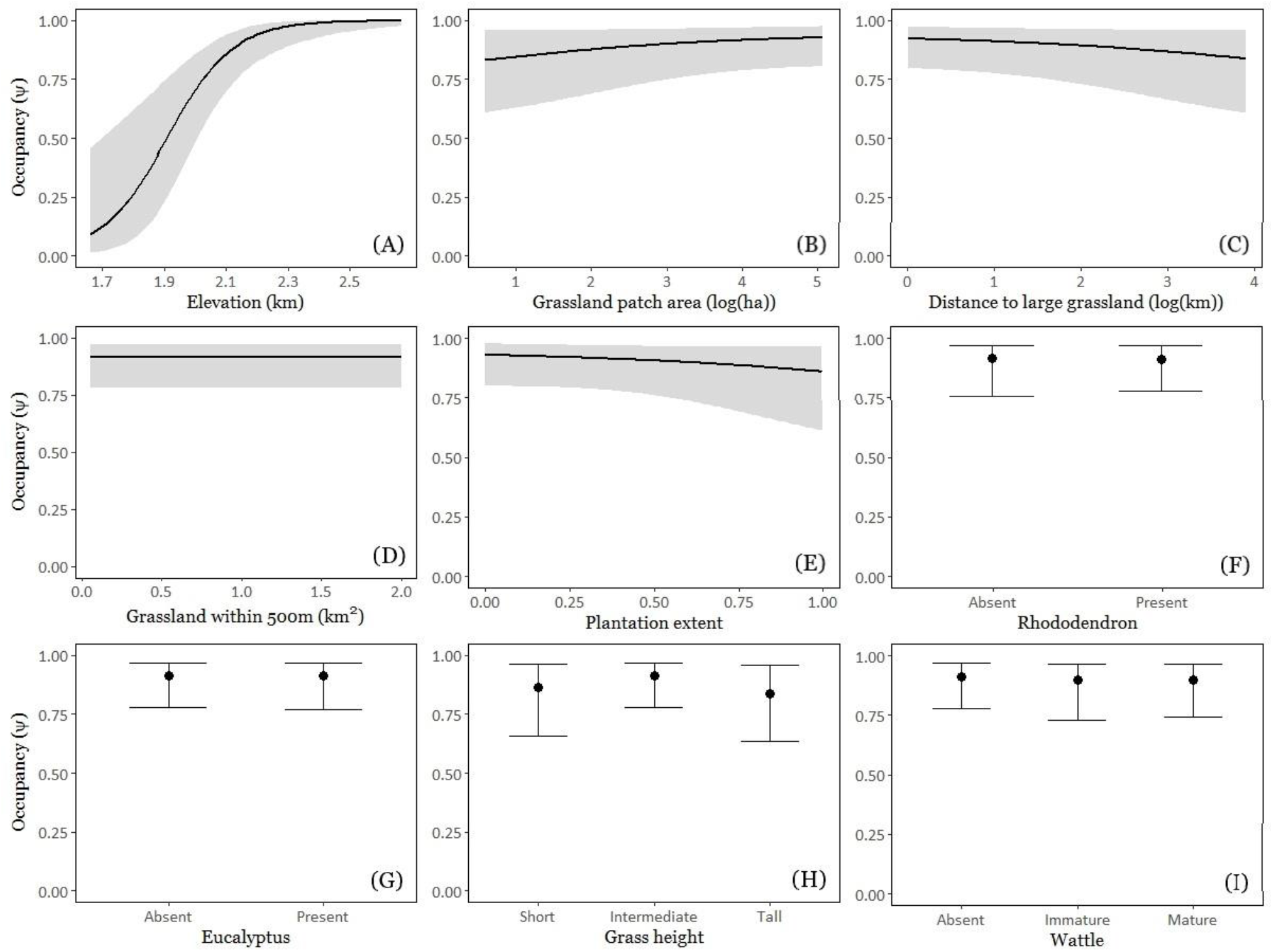
A-I: Model-averaged predicted occupancy of the Nilgiri pipit in response to all covariates appearing in models; maximum elevation within a site (A), log grassland patch area (B), log distance to the nearest grassland larger than 1.5 km^2^ (C), grassland within 500m of the site boundary (D), extent of plantation within the site (E), presence of rhododendron (F), presence of *Eucalyptus* (G), grass height (H), and wattle maturity (I). All other variables are set to mean or modal values for continuous and categorical covariates, respectively. Probability of occupancy is plotted over the observed range of values of each predictor.

